# BehaviorDEPOT: a tool for automated behavior classification and analysis in rodents

**DOI:** 10.1101/2021.06.20.449150

**Authors:** Christopher J Gabriel, Zachary Zeidler, Benita Jin, Changliang Guo, Anna Wu, Molly Delaney, Jovian Cheung, Lauren E. DiFazio, Melissa J. Sharpe, Daniel Aharoni, Scott A. Wilke, Laura A. DeNardo

## Abstract

Quantitative descriptions of animal behavior are essential to understand the underlying neural substrates. Many behavioral analyses are performed by hand or with expensive and inflexible commercial software that often fail on animals with attached head implants, such as those used for *in vivo* optogenetics and calcium imaging. With the development of machine learning algorithms that can estimate animal positions across time and space, it is becoming easier for users with no prior coding experience to perform automated animal tracking in behavioral video recordings. Yet classifying discrete behaviors based on positional tracking data remains a significant challenge. To achieve this, we must start with reliable ground truth definitions of behavior, a process that is hindered by unreliable human annotations. To overcome these barriers, we developed BehaviorDEPOT (DEcoding behavior based on POsitional Tracking), a MATLAB-based application comprising six independent modules and a graphical user interface. In the Analysis Module we provide hard-coded classifiers for freezing and rearing. Optionally applied spatiotemporal filters allow users to analyze behaviors in varied experimental designs (e.g. cued tasks or optogenetic manipulations). Even inexperienced users can generate organized behavioral data arrays that can be seamlessly aligned with neurophysiological recordings for detailed analyses of the neural substrates. Four additional modules create an easy-to-use pipeline for establishing reliable ground-truth definitions of behaviors as well as custom behavioral classifiers. Finally, our Experiment Module runs fear conditioning experiments using an Arduino-based design that interfaces with commercial hardware and significantly reduces associated costs. We demonstrate the utility and flexibility of BehaviorDEPOT in widely used behavioral assays including fear conditioning, avoidance, and decision-making tasks. We also demonstrate the robustness of the BehaviorDEPOT freezing classifier across multiple camera types and in mice and rats wearing optogenetic patch cables and head-mounted Miniscopes. BehaviorDEPOT provides a simple, flexible, automated pipeline to move from pose tracking to reliably quantifying a wide variety of task-relevant behaviors.

## Introduction

A central goal of neuroscience is to discover relationships between neural activity and behavior. Discrete rodent behaviors represent outward manifestations of cognitive and emotional processes. It is critical to classify such behaviors rapidly and reliably with high spatiotemporal precision. The recent explosion of techniques for manipulating or recording activity in neural circuits is advancing behavioral neuroscience at an unprecedented pace^1^. However, a major bottleneck has been aligning these data with coincident behavior, especially for laboratories without established expertise in this area. Much of this work has been done manually, but automated detection of freely moving animal behaviors is faster, expands the parameter space that can be explored, and removes human biases. The standardization promised by such methods also enhances the rigor and reproducibility of results across research groups. However, these analyses can be technically challenging for researchers to develop in-house.

Recently, behavioral neuroscientists have embraced machine learning algorithms for tracking animal positions across time and space^2^. Free and easy-to-use software packages such as DeepLabCut (**DLC**) allow even inexperienced users to train deep neural networks that automatically track user-defined points on an animal’s body^3^. The static arrangement of those points at any moment represents a pose. The challenge then is to classify temporal pose sequences as discrete behaviors that represent useful information about the cognitive or emotional state of the animal. This is a major challenge for two main reasons: 1) developing automated classifiers typically requires advanced programming skills and 2) developing robust classifiers requires accurate ground-truth definitions of behaviors, but human annotations are error-prone and unreliable.

Quantifying animal behavior can be complex and technically challenging. Some behaviors can be defined exclusively by the animal’s pose, such as during freezing in response to fearful stimuli. Other common assays, like the T-maze, elevated plus maze, and open field test, utilize spatiotemporal features to define behavior. Many labs currently score behaviors manually, a time-consuming, laborious, and error-prone process. Others use commercially available software that can be prohibitively expensive and fail when used with common headgear for *in vivo* neurophysiology. Moreover, manual scoring and commercial software typically only report when behavior was present, providing a limited view of the animal’s behavior. Robustly analyzing spatial and temporal information can enhance behavioral assessment by providing important layers of information about the animal’s emotional and physical status.

To overcome these challenges, we developed an open-source software package for behavior **DE**coding based on **PO**sitional **T**racking (BehaviorDEPOT) that is robust, flexible and easy to use. BehaviorDEPOT is a MATLAB-based application comprising six independent modules (Figure 1). With its graphical user interface (GUI), no coding experience is required to use BehaviorDEPOT, but experienced users can tailor the software to their liking. The Analysis Module imports behavior videos and accompanying pose tracking data and classifies behavior on a frame-by-frame basis. It includes hard-coded classifiers for freezing, rearing, moving, and jumping. With optionally applied spatiotemporal filters, it can perform automated behavioral analysis for any experimental design (e.g., cued tasks or optogenetics experiments). Vectorized data is saved in MATLAB data arrays that facilitate additional user-defined automated analyses. The Analysis Module also features customizable visualization tools that integrate behavioral classification with spatiotemporal trajectory mapping. The Experiment Module can run fear conditioning experiments using an Arduino-based protocol that is compatible with commercially available shockers, arenas, and lasers for optogenetics.

**Figure 1.**
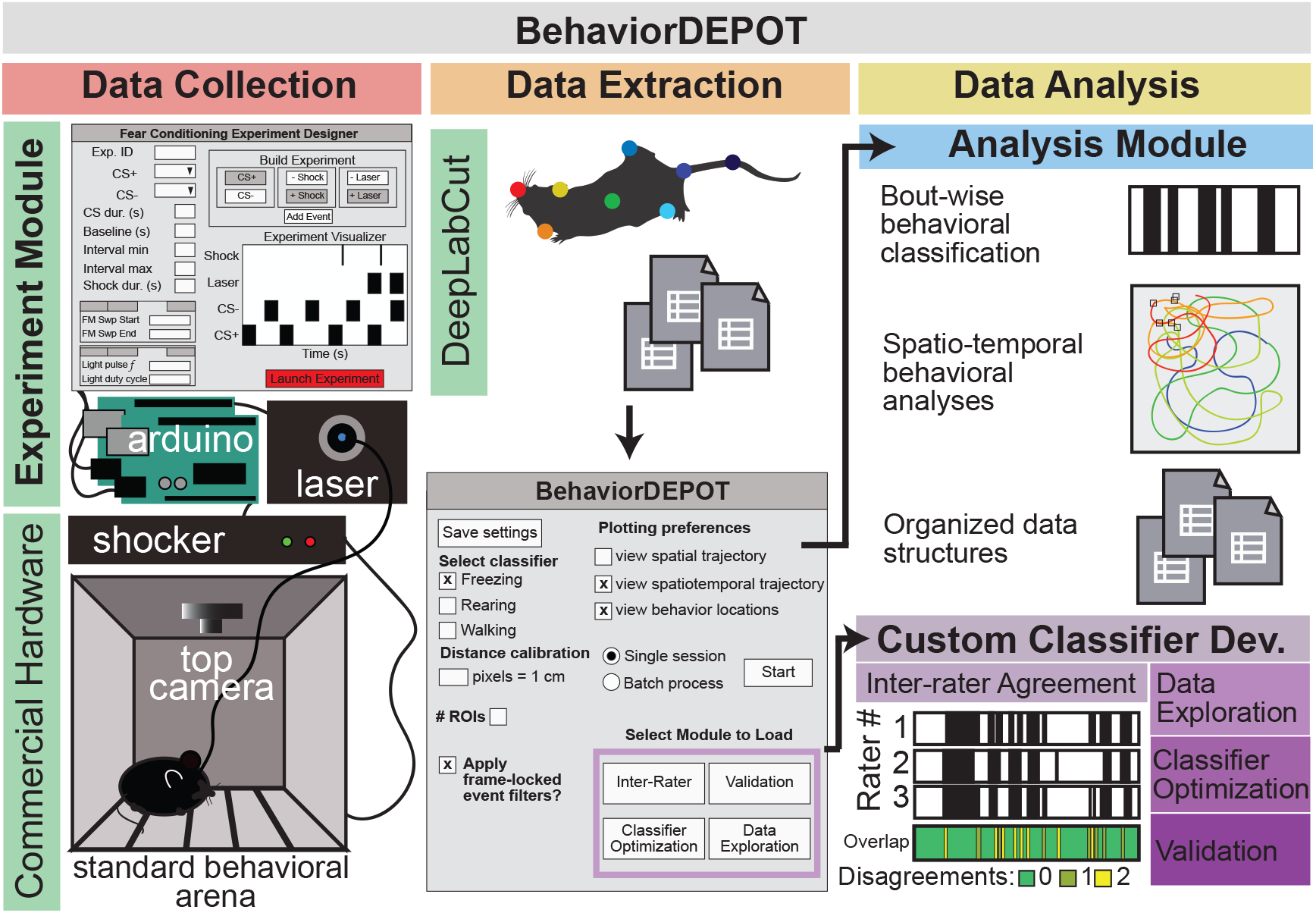
BehaviorDEPOT overview. The Experiment Module is a MATLAB GUI based application that allows users to design and run fear conditioning experiments. The software uses Arduinos to interface with commercially available hardware (e.g., shockers and lasers) to control stimuli. Behavior videos are acquired with webcams or machine-learning quality cameras. Video recordings are analyzed with pre-trained DeepLabCut models. The video and pose estimates are the inputs for a second MATLAB GUI that controls the Analysis Module for customizable analyses of freezing, rearing, and moving. Four additional modules help users develop custom classifiers.

Four additional independent modules guide users through designing custom classifiers in a manner that requires no programming experience. A major hurdle in developing classifiers is settling on a ground truth definition of the behavior of interest. It is difficult, particularly for unskilled raters, to annotate hours of behavior videos without introducing errors or having annotations diverge amongst individuals and across labs. However, several skilled raters, focusing on establishing ground-truth definitions can establish behaviors which can then be more reliably annotated by automated classifiers. The Inter-Rater Module simplifies this process by performing automated comparisons of manual annotations from multiple human raters. Taking in frame-by-frame human annotations, it calculates the level of inter-rater agreement and highlights points of disagreement so the raters can refine their annotations until they converge maximally. The Inter-Rater Module can function as an independent unit and can thus support classifier development with any application. The additional modules help users perform parameter exploration, optimization, and validation of new classifiers (Figure 1).

Our goal with BehaviorDEPOT was to create a free and user-friendly tool for automated classification and spatiotemporal analysis of commonly studied behaviors. We demonstrate BehaviorDEPOT’s utility with three frequently used behavioral assays, yet it is highly customizable, assisting users to generate new classifiers with minimal coding involved. Other open-source software packages that transform animal poses into discrete behaviors focus on social behaviors or behavioral syllables that are not necessarily associated with specific emotional or motivational states (MARS^4^, SimBA^5^, MoSeq^6^, B-SOiD^7^, and DANNCE^8^). To our knowledge, BehaviorDEPOT provides the only open-source pose tracking-based freezing classifier that operates via a user-friendly GUI. Compared to existing open-source freezing detection software^9^, BehaviorDEPOT can both implement and analyze fear conditioning experiments. BehaviorDEPOT can perform batched data analyses and has the unique ability to integrate behavior classification with the spatiotemporal trajectory of the animal. This facilitates analysis of assays in which the experimenter may want to quantify not only how much an animal performs a particular behavior overall, but where and when it does so.

## Results

BehaviorDEPOT functions as a MATLAB application or as a standalone executable file. Through its GUI, users can interact with six independent modules (Figure 1, Figure S1). The Analysis Module smooths pose tracking data and quantifies freezing in addition to a variety of defensive and ambulatory behaviors including escape, running, and walking. Users can implement custom temporal and spatial filters to analyze behaviors in varied experimental designs. The other four modules allow users to create and optimize custom classifiers with little-to-no programming experience.

### Development of the BehaviorDEPOT Freezing Classifier

BehaviorDEPOT combines a convolution-based smoothing operation with a low-pass filter to identify periods of freezing (Figure 2A–D), defined as the absence of movement except for respiration. We began by training two different deep neural networks in DeepLabCut. One is based on videos recorded with a machine-learning quality camera at 50 frames per second (fps). This network tracks 8 points on the body (Figure 2A). The second network is based on videos recorded with a standard webcam at 30fps. On the webcam, lower sensor quality and lower frame rate produces more blur in the recorded images, so we only tracked 4 body parts (nose, ears, tail-base; Figure 2). While it is possible to generate a single DLC network that can estimate the positions of body parts with reasonable precision in both video types, we found that more tailored networks produced lower jitter (i.e., small changes in point placement from frame to frame) around the estimated points. Low jitter proved to be critical for creating the most robust freezing classifier. Users can select the classifier that works best for their videos.

**Figure 2.**
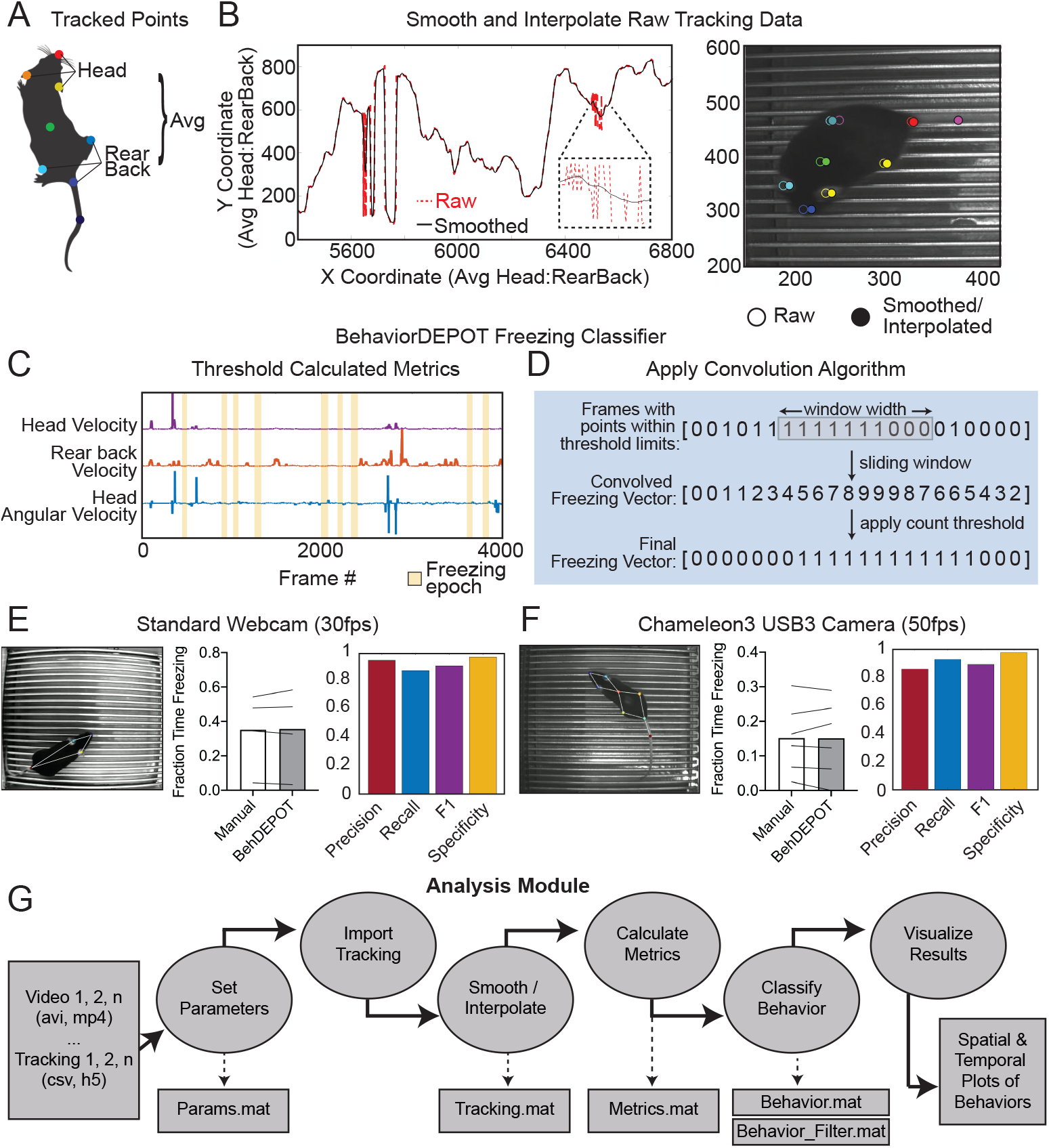
The Analysis Module. A. Metrics based on individual tracked points and weighted averages are calculated and stored in BehaviorDEPOT data matrices. B. Visualization of the effects of the LOWESS smoothing and interpolation algorithms for the weighted average of head and rear back (left) and for all tracked points in a representative example frame (right). C. Visualization of metrics that form the basis of the BehaviorDEPOT freezing classifier. Colored lines represent framewise calculated values for each metric. Yellow bars indicate freezing epochs. D. Visualization of the convolution algorithm employed by the BehaviorDEPOT freezing classifier. A sliding window of a specified width produces a convolved freezing vector in which each value represents the number of freezing frames visible in the window at a given frame. An adjustable count threshold converts the convolved freezing vector into the final binary freezing vector. E. Evaluation of freezing classifier performance on videos recorded at 30fps with a standard webcam (Precision: 0.95, Recall: 0.88, F1: 0.97, Specificity: 0.91). F. Evaluation of freezing classifier performance on videos recorded at 50 fps with a machine learning-quality camera (Precision: 0.86, Recall: 0.92, F1: 0.89, Specificity: 0.97). G. The Analysis Module workflow. Videos and accompanying pose tracking data are the inputs. Pose tracking and behavioral data is vectorized and saved in MATLAB structures to facilitate subsequent analyses.

DLC produces a list of comma-separated values that contains framewise estimates of the x-y coordinates for designated body parts as well as a likelihood statistic for each estimated point. We first applied a threshold based on DLC likelihood (p < 0.1) and performed a Hampel transformation^10^ to remove outliers. Then, a LOWESS, local regression smoothing method was applied to the data^11^, and sub-threshold tracking values were estimated using surrounding data and spline interpolation (Figure 2B). We calculated statistics based on tracked points including linear velocity, acceleration, head angle, and angular velocity. After exploring the parameter space, we empirically determined that thresholding the velocity of a weighted average of 3–6 body parts (depending on the frame rate of the video recording) and the angle of head movements produced the best-performing freezing classifier (Figure 2C). We applied a sliding window to produce a convolved freezing vector in which each value represents the number of freezing frames visible in the window at a given frame. We then applied an adjustable count threshold to convert the convolved freezing vector into the final binary freezing vector (Figure 2D).

We validated the performance of the freezing classifier by comparing it to manual scoring by three trained raters. To ensure our classifiers were robust, our validation data sets included ~30,000 frames from 4–6 videos (at least 6000 frames per video) that were recorded in distinct behavioral chambers under varied lighting conditions. Human ratings were indistinguishable from classifier performance (Figure 2E,F). Accuracy of the freezing classifier was estimated based on classifier precision (number of true positive frames / sum of true positive and false positive frames), recall (number of true positive frames / sum of true positive and false negative frames), F1 score (two times the product of precision and recall / sum of precision and recall) and specificity (number of true negative frames / sum of true negative and false positive frames). Precision and recall quantify the positive predictive value of the classifier against the tendency to produce false positive or false negative errors, respectively. The F1 score, the harmonic mean of the precision and recall, is useful as a summary statistic of overall classifier performance. Specificity quantifies the classifier’s ability to accurately label true negative values and helps ensure that the classifier is capturing data from only a single annotated behavior. The high values in all categories (Figure 2E,F) indicated that freezing classifiers based on webcams and machine learning quality cameras both had excellent performance.

### The Analysis Module

The Analysis Module imports videos and accompanying pose tracking data, smooths the data using a user-specified method, and calculates movement statistics based on tracked points (e.g., linear velocity, angular velocity, acceleration). It then analyzes behavior in a framewise manner by implementing custom classifiers that threshold movement and position statistics in unique combinations. In the GUI, users indicate if they want to analyze behaviors during particular time windows or within regions of interest (ROI), and whether they plan to do batched analyses. Depending on their choices, users are led through a series of windows in which they indicate the body parts that they tracked, ROI location(s), and when time windows of interest occurred. The Analysis Module analyzes behavior according to the user-defined specifications and saves the data in a set of MATLAB structures. The ‘Params’ structure stores the parameters of video recordings, smoothing, and arena metrics (e.g., ROI size and location). ‘Tracking’ organizes and stores pose data from DLC. ‘Metrics’ stores calculated movement and position statistics based on tracked points including linear velocity, angular velocity, and acceleration (Figure 2G). ‘Behavior’ stores bout-wise and vectorized representations of classified behaviors both in and out of user-defined spatiotemporal filters. Organizing the data this way facilitates additional automated analyses of the user’s choosing. Users can track any points they like, though they may have to adjust and validate the behavior thresholds using our Optimization and Validation Modules, described in upcoming sections.

The Analysis Module also generates a series of graphical data representations. Trajectory maps show when an animal was in a particular location and where behaviors occurred. Bout maps indicate when behaviors occurred and for how long. Additional graphs tell the user how much a particular behavior occurred during presentations of a cue or optogenetic stimulation period. These visual representations help users understand behavioral phenotypes in great spatiotemporal detail and can inform development of further custom analyses using the data arrays that the Analysis Module generates.

### Use Case 1: Optogenetics

In commercial freezing detection software, algorithms often fail when a rodent is wearing a patch cord for routine optogenetics experiments. We set out to validate BehaviorDEPOT’s utility in an optogenetics experiment. The medial prefrontal cortex (mPFC) plays a well-established role in fear memory retrieval^12,13^, extinction^14,15^, and generalization^14,16,17^. We performed an experiment to examine the role of mPFC in contextual fear memory generalization. While silencing mPFC subregions can promote fear memory generalization in remote memory^16,18^, less is known about its role in recent memory generalization. We used an adeno-associated virus (AAV) to express the soma-targeted neuronal silencer stGtACR2^19^ bilaterally in the mPFC and implanted optogenetic cannula directly above the AAV injection sites (Figure 3A, Figure S2). We performed contextual fear conditioning (CFC) and then tested animals’ fear memory 24 hours later by recording their behavior in both the fearful context and in a novel environment that was not previously paired with shocks (Figure 3B). We estimated the animals’ position on each frame using an optogenetics-specific DLC network that tracked 9 points on the animal, including the fiber-optic cannula (Figure 3C). We then analyzed freezing levels with BehaviorDEPOT. Fear conditioned mice readily froze following shocks during CFC, while non-shocked controls did not (Figure 3D). Silencing mPFC in previously shocked animals significantly enhanced freezing in the novel context but did not affect freezing in the fearful context (Figure 3E,F). mPFC silencing thus produced a significant decrease in the discrimination index in fear conditioned mice (Figure 3G), indicating that mPFC plays a key role in the specificity of recent fear memories. In all analyses, BehaviorDEPOT performance was indistinguishable from a highly trained human rater (Figure 3E–G) indicating that the combination of the optogenetics-specific DLC model and BehaviorDEPOT achieves robust and accurate freezing detection.

**Figure 3.**
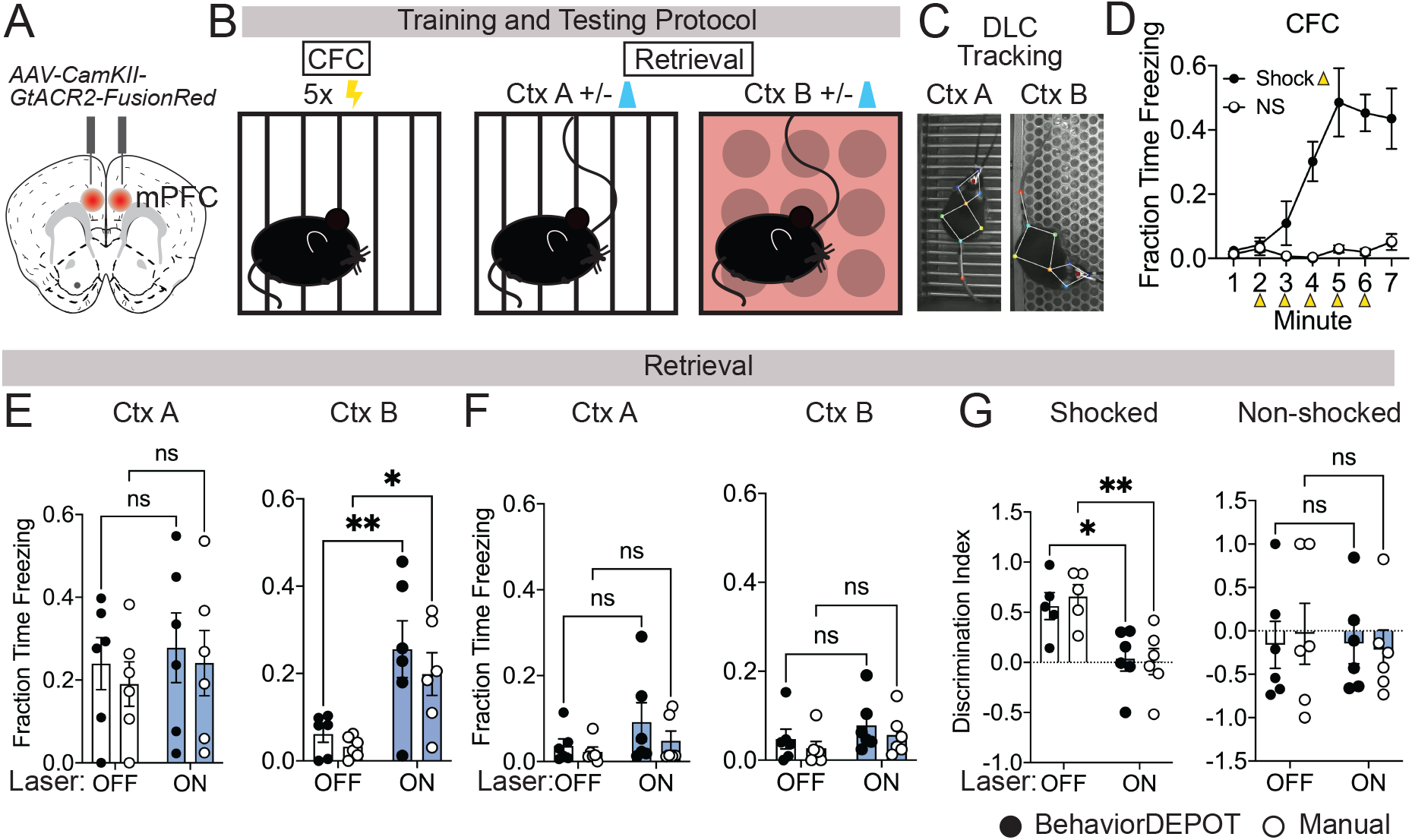
Use Case 1: Optogenetics. A. AAV1-CamKII-GtACR2-FusionRed was injected bilaterally into medial prefrontal cortex (mPFC). B. Behavioral protocol. Mice underwent contextual fear conditioning on day 1. On day 2, mice were returned to the conditioned context or a novel context in a counterbalanced fashion and received 2×2 min 473nm laser stimulation separated by 2 min laser off intervals. C. Example DLC tracking of mice attached to patch cords in different contexts. D. Quantification of contextual freezing during training analyzed with BehaviorDEPOT. E–F. Comparing human annotations to BehaviorDEPOT freezing classifier. E. Shocked mice: freezing in context A (left) and context B (right) with and without mPFC silencing (CtxA: *F*_*laser*_(1,10)=0.42, *P*=0.53; *F*_*rater*_(1,10)=0.35, *P*=0.57; CtxB: *F*_*laser*_(1,10)=26.51, *P*=0.0004; *F*_*rater*_(1,10)=0.08, *P*=0.78; Two-way repeated measures ANOVA, N=6 mice per group). F. Non-shocked controls: freezing in context A (left) and context B (right) with and without mPFC silencing (*F*_laser_(1,10)=3.60, *P*=0.09; *F*_rater_(1,10)=0.79, *P*=0.39; Two-way repeated measures ANOVA, N=6 mice per group). G. Discrimination index = (FreezeA - FreezeB) / (FreezeA + FreezeB) for shocked mice (*F*_laser_(1,10)=17.54, *P*=0.002; *F*_rater_(1,8)=0.09, *P*=0.77; Mixed-effects analysis, N=5–6 per group) and non-shocked controls (*F*_laser_(1,10)=0.07, *P*=0.80; *F*_rater_(1,8)=0.02, *P*=0.90; Two-way ANOVA). Error bars represent S.E.M.

### Use Case 2: Ca^2+^ imaging with Miniscopes during signaled avoidance

As new open-source tools for neurophysiology become available, increasing numbers of labs are performing simultaneous neurophysiological and behavioral recordings to understand the neural mechanisms of complex behaviors. Recent advances in miniature head-mounted microscopes now allow us to image the activity of hundreds of neurons simultaneously in freely moving animals^20–22^. These miniscopes pair with genetically encoded Ca^2+^ indicators^23^ that can be targeted to genetically, anatomically, or behaviorally-defined neuronal populations, and GRIN lenses^24^ that can be targeted anywhere in the brain. With these tools in hand, we are now able to discover how the encoding of complex cognitive and emotional behaviors maps onto specific cell types across the brain. By recording the activity of hundreds of neurons simultaneously, we can also begin to understand the population codes that produce complex, freely moving behaviors^25,26^. To do so, however, we need improved methods that allow us to rapidly and accurately quantify freely moving behaviors with reference to salient environmental stimuli and to align these detailed behavioral measurements with neurophysiological recordings. BehaviorDEPOT directly addresses these deficits by automatically integrating and quantifying spatial information (e.g animal location), temporal information (e.g. onset of conditioned cues), and the time and location at which discrete behaviors occurred. Stored in vectorized data structures in a framewise manner, these detailed behavioral data can be instantly aligned with physiological recordings.

Here we demonstrate the utility of BehaviorDEPOT for aligning behavioral measurements with Ca^2+^ signals during a signaled avoidance task that has temporally and spatially salient features. In our task, which is a refinement of platform mediated avoidance^27^, a fear conditioned tone prompts mice to navigate to a safety platform that protects them from receiving a footshock. During a behavioral session, animals toggle between conditioned freezing and environmental exploration in a manner that is modulated by temporal (e.g. conditioned tones) and spatial (e.g. safety platform) factors. In this task, the medial prefrontal cortex acts through downstream target regions to determine whether an animal approaches or avoids environmental stimuli^28^. To determine how ensembles of prefrontal neurons drive behavior during PMA, we recorded the activity of hundreds of mPFC neurons in freely behaving animals using head-mounted microendoscopes (UCLA Miniscopes^21,22^) while simultaneously recording behavior using a new open-source USB camera, the UCLA MiniCAM.

The MiniCAM is an open-source behavioral imaging platform that natively integrates and synchronizes with open-source UCLA Miniscope hardware and software (Figure 4A). It is composed of an M12 optical lens mount, a custom printed circuit board housing a CMOS image senor and supporting electronics, an LED illumination ring, and a 3D printed case. The MiniCAM is powered and communicates over a single coaxial cable that can be up to 15 meters long. The coaxial cable connects to a Miniscope data acquisition board (DAQ) which then connects over USB3 to a host computer. A range of commercial M12 lenses can be used to select the view angle of the camera system. The image sensor used is a 5MP CMOS image sensor (MT9P031I12STM-DP, ON Semiconductor) with 2592 x 1944 pixel resolution and a full resolution frame rate of approximately 14FPS. For this application, the MiniCAM’s pixels were binned and cropped to achieve 1024×768 pixels at approximately 50FPS. The optional LED illumination ring uses 16 adjustable red LEDs (LTST-C190KRKT, Lite-On Inc., 639nm peak wavelength) for illumination in dark environments (Figure 4A). We trained a separate DLC network for videos of animals wearing Miniscopes recorded with MiniCAMs. Our network tracked 8 points on the body (ears, nose, midback, hips, tailbase, and tail) and the Miniscope itself (Figure 4B). Using the output of this network, BehaviorDEPOT classified freezing with high fidelity in a manner that was indistinguishable from highly trained users (Figure 4C).

**Figure 4.**
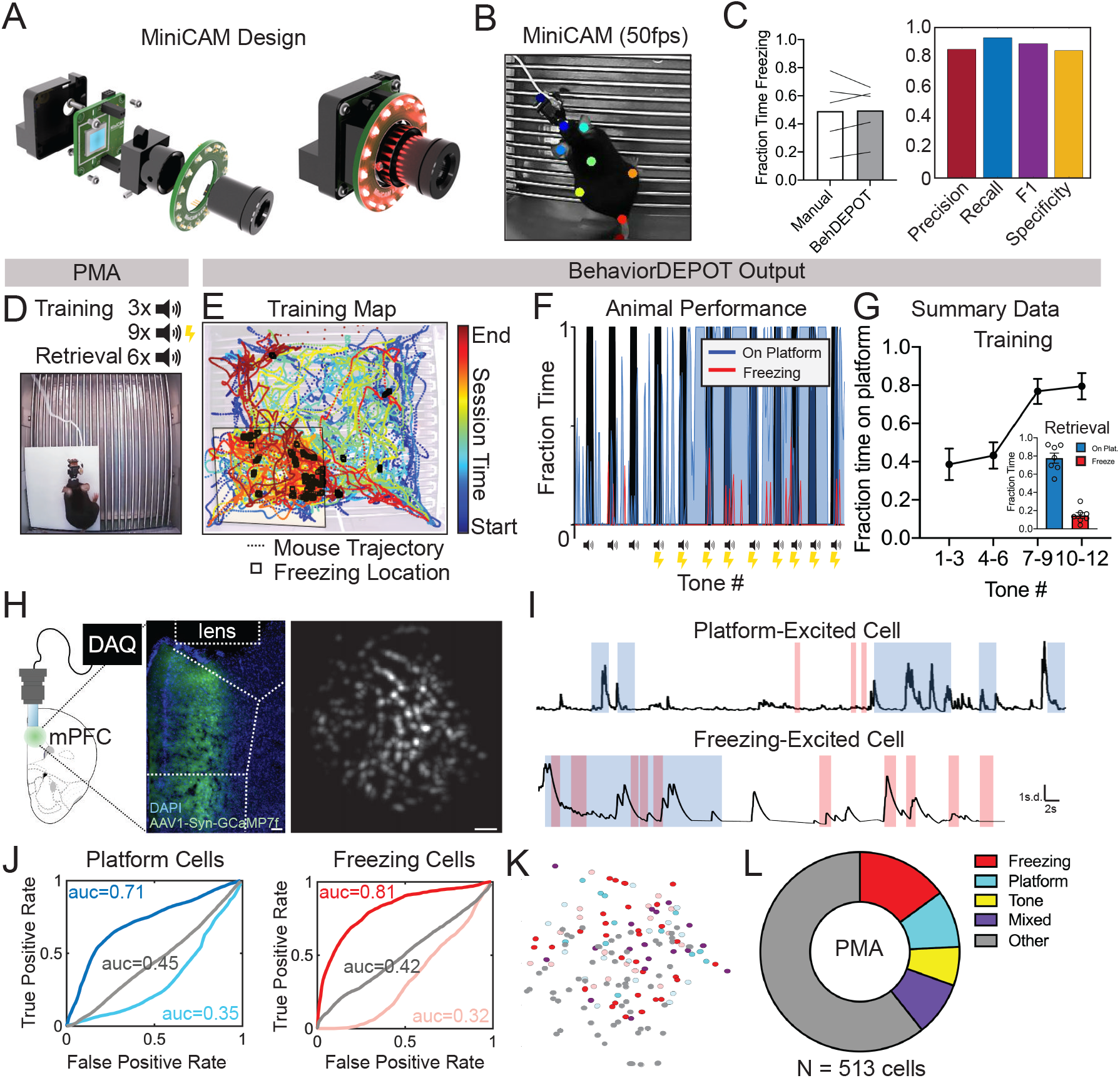
Use Case 2: Mice wearing Miniscopes. A. Design for MiniCAM, an open-source camera designed to interface with Miniscopes and pose tracking. B. Still frame from MiniCAM recording of mouse wearing a V4 Minscope. DLC tracked points are labeled with rainbow dots. C. Performance of freezing classifier on videos of mouse wearing Miniscope recorded with MiniCAM (Precision: 0.85; Recall: 0.93; F1 Score: 0.89; Specificity: 0.84). D. Task design. E. Sample BehaviorDEPOT output for mouse wearing Miniscope during PMA. Map displays animal position over time as well as freezing locations (black squares). F. Temporal alignment of time on the platform (blue), time freezing (black), and tones. G. Summary data for training and retrieval. H. GCaMP7-exressing mPFC neurons imaged through a V4 Miniscope. I. Example Ca^2+^ traces from platform (top) and tone (bottom) modulated cells during time on the platform (blue) or time freezing (pink). J. Receiver operating characteristic (ROC) curves that were calculated for platform-modulated cells (excited cell: auc-0.71; suppressed cell: auc=0.35, unmodulated cell: auc=0.45) and freezing-modulated cells (excited cell: auc=0.81; suppressed cell: auc=0.32; unmodulated cell: auc=0.42). K. Example field of view showing locations of freezing- and platform-modulated mPFC neurons. L. Proportion of modulated cells of each functional type from 513 cells recorded across 3 mice. Scale bars, 100um. Error bars represent S.E.M.

We used BehaviorDEPOT to analyze behavior during PMA so we could align it to the underlying neural activity. In this task, animals hear 3 baseline tones (4kHz, 30s) followed by 9 tones that co-terminate with a mild foot shock (0.15mA, 2s). An acrylic platform occupies 25% of the electrified grid floor, providing a safe place to avoid the shock. The following day, animals are presented with 6 unreinforced tones, and we measure the resulting avoidance and freezing responses (Figure 4D). The Analysis Module automatically produces maps that make it quick and easy to assess the spatiotemporal characteristics of rodent behavior. In our representative example, the color-coded trajectory and freezing locations (denoted as black squares) converge on the platform at the end of the session, indicating the mouse indeed learned to avoid shocks by entering the platform (Figure 4E). The vectorized session data from the Experiment Module (e.g. tone times, shock times) and behavioral data from the Analysis module can be easily integrated for additional analyses. We visualized the temporal dynamics of learning by overlaying fraction time on the platform and freezing on the tone periods (Figure 4F). We also used BehaviorDEPOT to produce summary data, showing that mice readily learned and remembered the cue-avoidance association as expected (Figure 4G).

During a retrieval session, we performed simultaneous recordings of neural activity using a UCLA Miniscope and behavior using a UCLA MiniCAM (Figure 4H). Using MIN1PIPE^29^, we extracted and processed neural signals from 513 mPFC neurons across 3 mice. We then determined whether individual neurons encoded specific behaviors that we had quantified using BehaviorDEPOT. We computed a receiver operator characteristic (ROC) curve that measures a neuron’s stimulus detection strength over a range of thresholds (Figure 4I,J). We identified numerous neurons that were modulated by freezing and avoidance on the safety platform. These neurons were organized in a salt and pepper manner in mPFC (Figure 4K). Nearly half of all neurons that exhibited task relevant behavior and were specific for either freezing or avoiding on the platform, or the combination of both (Figure 4L).

### The Inter-Rater Module

A major challenge in behavioral neuroscience is establishing reliable ground truth definitions of behavior. Even when using automated behavioral classification systems like BehaviorDEPOT, it is critical to have reliable behavioral annotations from multiple human raters to validate the performance of the classifier. The Inter-Rater Module helps users establish reliable ground truth behavioral definitions in two ways. First, the Inter-Rater Module imports manual behavioral scoring from human raters and automatically compares the scores on a frame-by-frame basis. It calculates the level of agreement and reports points of disagreement both numerically and graphically. In response to the initial output of the Inter-Rater Module, human raters can discuss the points of disagreement and modify their manual ratings accordingly. The automated features of the Inter-Rater Module make it fast and easy to perform iterative comparisons of manual annotations, interleaved with human updates, until a satisfactory level of agreement is achieved.

The Inter-Rater Module prompts the user to select a file directory containing one or more human annotation files and automatically imports the data into MATLAB. The user then selects the annotations to include in the analysis, the reference dataset, and the behavior of interest. After all parameters are set, the module compares each set of annotations to the reference, scoring the annotations frame-by-frame as true positive, true negative, false positive, or false negative for each rater. These values are first used to calculate percent overlap and percent error between all raters. Precision, recall, specificity, and F1 score are calculated and reported for each rater relative to the chosen reference. Additionally, visualizations of frame-by-frame percent agreement and user-by-user disagreement are automatically generated to assist identifying areas of conflict between users (Figure 5A). We developed the BehaviorDEPOT freezing classifier based on the averaged ratings of three highly trained raters. Here, the Inter-Rater Module demonstrates visualizations of agreement levels for three highly trained raters and a novice freezing rater (Figure 5B). By illustrating frames with high level of disagreement, the Inter-Rater Module can help labs train new human annotators while building ground truth definitions for new classifiers.

**Figure 5.**
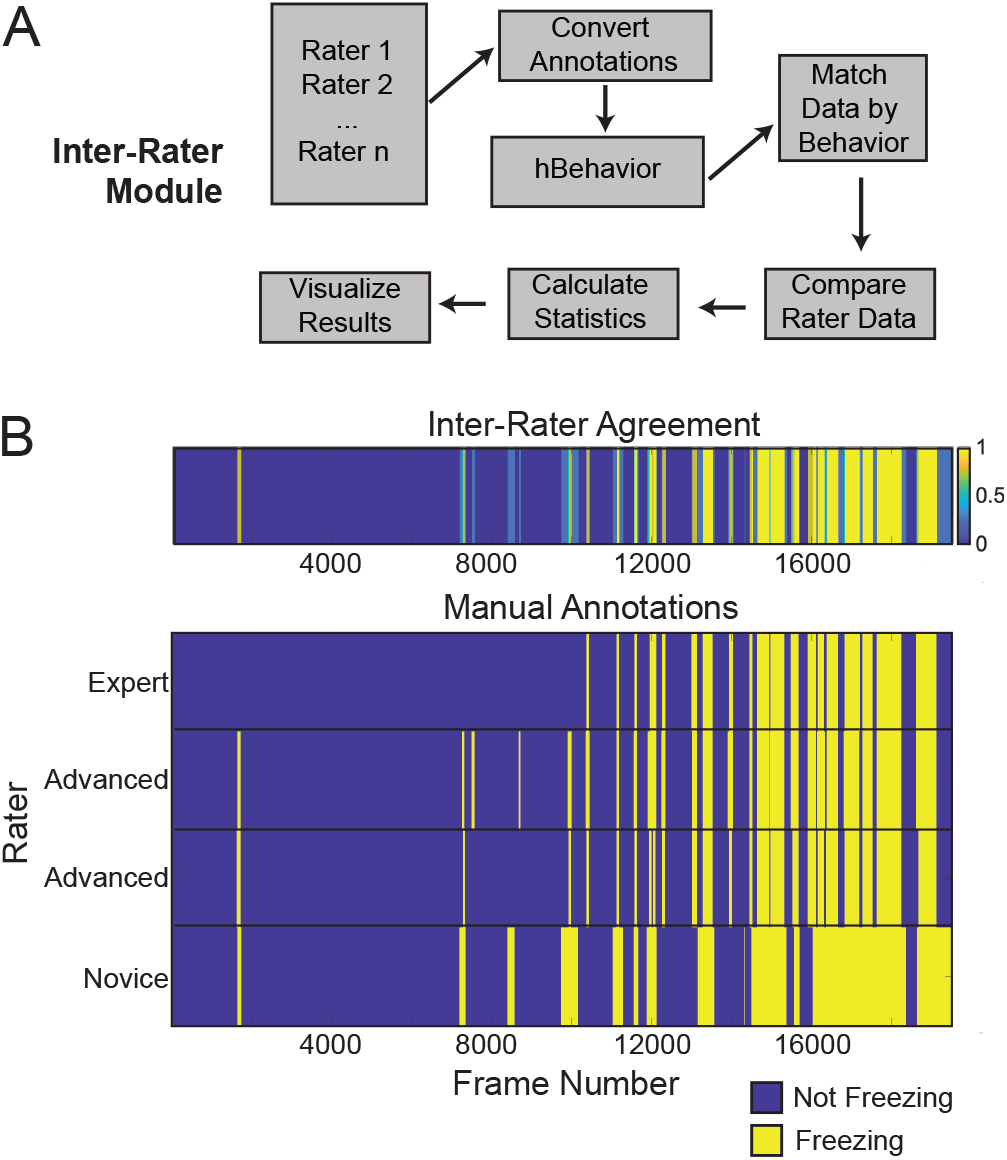
Inter-Rater Module. A. The Inter-Rater Module workflow. B. The Inter-Rater Module produces visualizations of framewise agreement levels (top) based on manual annotations from multiple human raters (bottom).

### The Data Exploration Module

The Data Exploration Module is designed to help refine existing classifiers and to generate entirely new ones. When developing behavior classifiers, it is advantageous to explore the parameter space of positional tracking data and identify the metrics that track most closely with a behavior of interest. The Data Exploration Module allows users to explore the metrics that BehaviorDEPOT calculates based on tracked points and determine whether chosen metrics separate behavior data accurately, using human labels as a reference. The chosen metrics are used to generate a generalized linear model (GLM) that estimates the likelihood that the behavior is present in a frame, given different values of the selected metrics (Figure 6A). This module allows users to easily identify and compare metrics that can form the basis of new classifiers or enhance the performance of existing ones.

**Figure 6.**
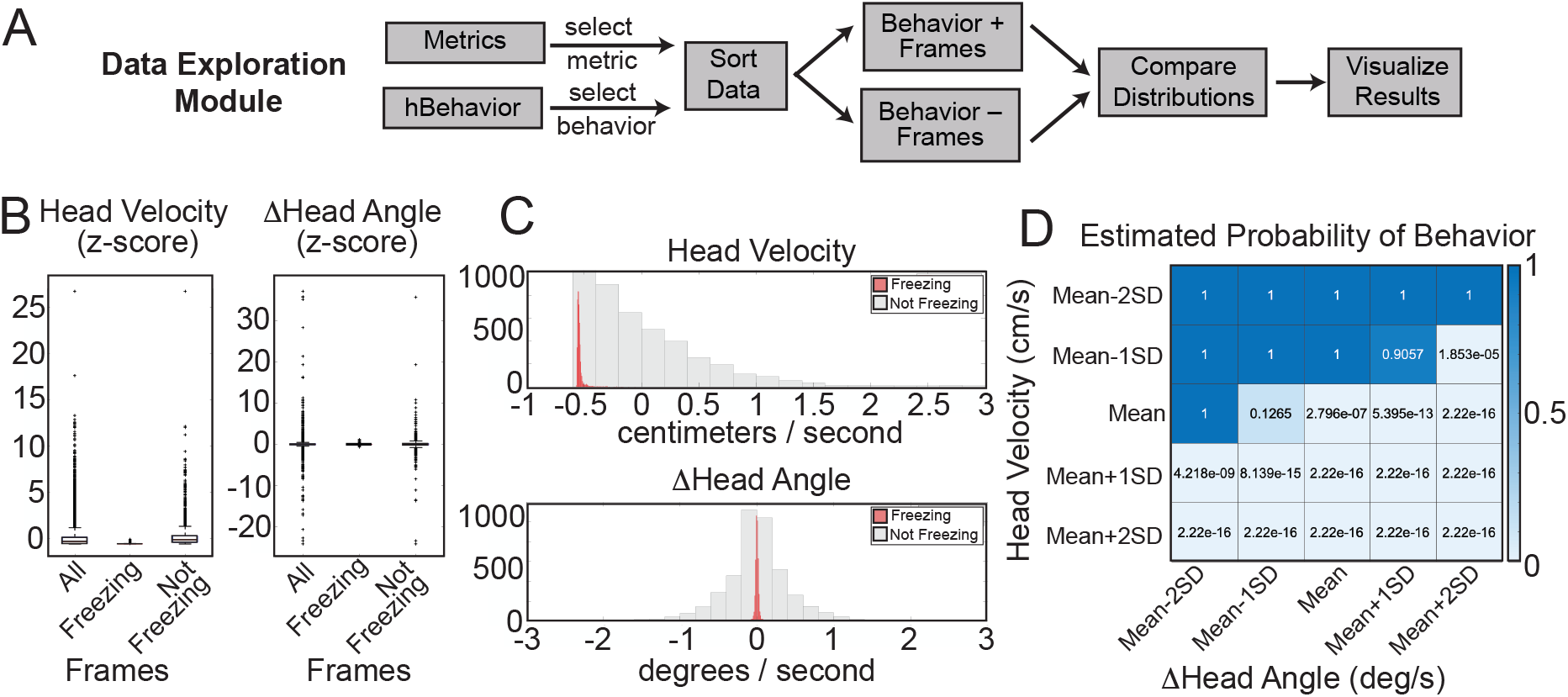
The Data Exploration Module. A. The Data Exploration Module takes in tracked metrics from the Analysis Module and the human annotations of behavior. It sorts the data in a framewise manner, separating frames containing the behavior of interest from those without and then visualizes and compares the distribution of values for a metric of interest. B. Distributions of Z-scored values for head velocity (left) and change in head angle (right) are distinct for freezing vs. not freezing frames. Box plots represent median, 25^th^, and 75^th^ percentile. Error bars extend to the most extreme point that is not an outlier C. Histograms showing distribution of values for head velocity (top) and change in head angle (bottom) for freezing (red) vs. not-freezing (grey) frames. D. A generalized linear model (GLM) computes the predictive power of given metrics for frames containing the behavior of interest.

The Data Exploration Module prompts the user to select a directory containing analyzed data and a human rater file, and the data is then automatically imported into MATLAB. Users can select two of the metrics from the analyzed file (e.g., head velocity, tail angular velocity, etc.) and a behavior label from the human annotations file. The module then creates two data distributions: one containing video frames labeled with the chosen behavior and a second consisting of the remaining video frames (Figure 6B). The module will randomly downsample the larger set to ensure that each distribution contains equal numbers of frames. The module then performs a series of analyses to quantify how reliably chosen metrics align with the behavior of interest. Histograms and boxplot representations of metric values for behavior-containing and non-behavior-containing frames help users identify metric values that reliably segregate with frames containing a behavior of interest (Figure 6C). Chosen metrics then form the basis of a GLM that predicts the likelihood that a frame captures the chosen behavior based on different values of each of the chosen metrics (Figure 6D). Metrics that are well-suited for behavior classification contrast with metrics on frames that do not contain the behavior and have a low standard deviation within the behavior set. Distributions of useful metrics also tend to differ substantially from the total set of frames, especially when compared to frames that do not contain the behavior. The GLM predictions are useful for determining which of the selected metrics best predict the behavior and whether they enhance the predictive value when combined.

### The Classifier Optimization Module

When developing a new classifier or optimizing our freezing classifier for use with externally generated data, BehaviorDEPOT’s Classifier Optimization Module allows user to rapidly assess how changes in classifier parameters (i.e. thresholded positional metrics) affect classifier performance (Figure 7A). To ensure that BehaviorDEPOT performs robustly across laboratories, we used the Classifier Optimization Module to adjust our freezing classifier for analysis of behavioral videos of rats and mice, respectively, that were recorded in two external laboratories using their own acquisition hardware and software (Figure 7B_1,2_). The Classifier Optimization Module takes in a BehaviorDEPOT-analyzed tracking file and associated human annotations and prompts the user to select up to two parameters from their chosen classifier. The user can then input a list of parameter values to test on the classifier. The module then runs the classifier using every combination of test values and calculates the performance (precision, recall, F1 score, and specificity) for each parameter combination. These values are stored in a MATLAB data array and automatically visualized using easy-to-read heatmaps. Importantly, the freezing classifier parameters that worked well for the data collected in our laboratory (change in head angle and head velocity) also worked well for the data collected in other labs. We used the Classifier Optimization Module to identify the combination of thresholds for two metrics (averaged head/rear back velocity and change in head angle, Figure 7C_1,2_) and the combination of window width and count threshold in the convolution algorithm (Figure 7D_1,2_) that produced maximal F1 scores. This module helps users optimize parameter values for individual classifiers and can be used in conjunction with the Data Exploration Module to build and test new behavior classifiers in a way that requires no prior coding experience.

**Figure 7.**
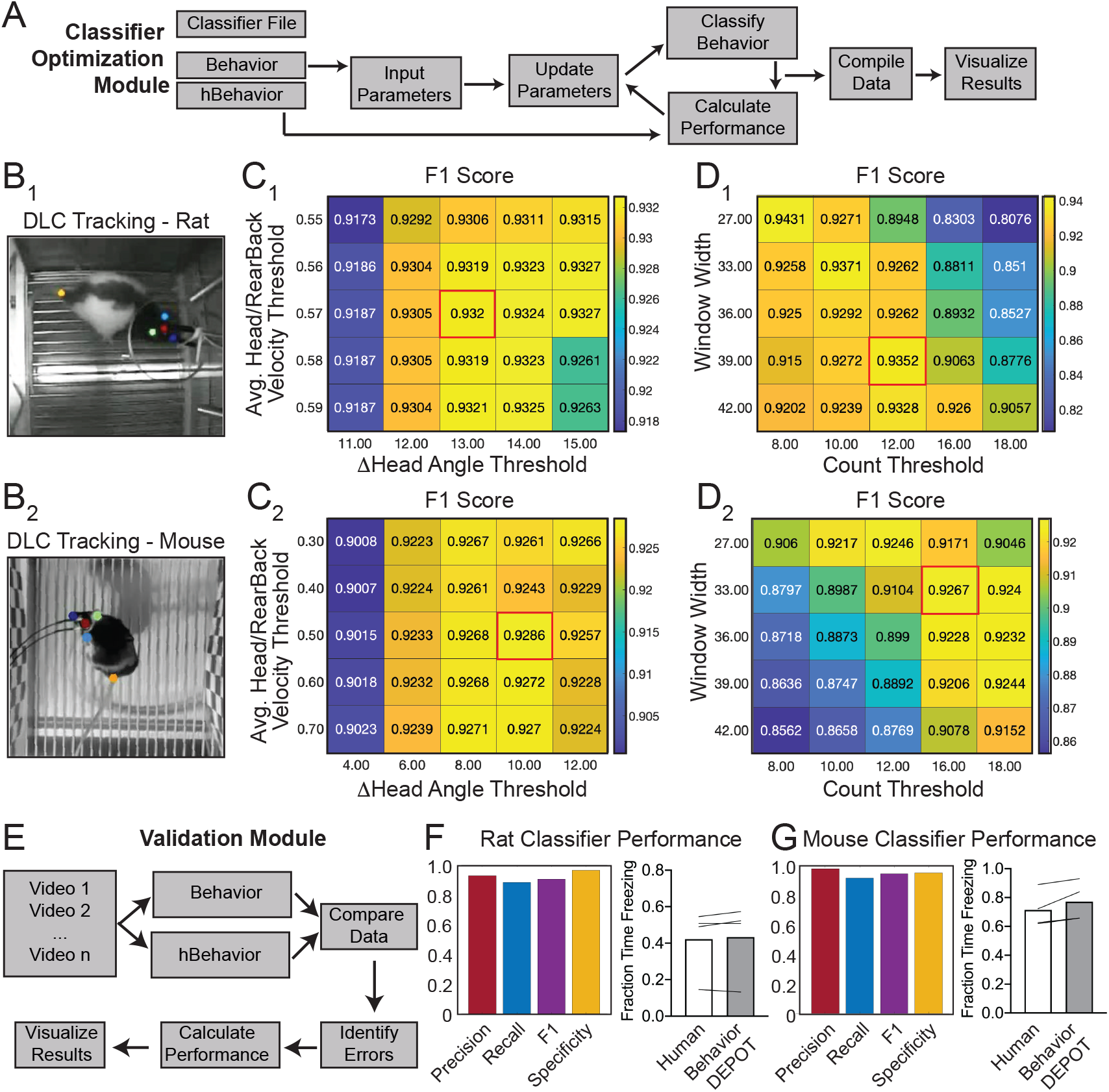
Use Case 3: Analysis of External Data using Classifier Optimization and Validation Modules. A. Classifier Optimization Module workflow. This module sweeps through a range of thresholds for statistics calculated based on tracked points and then compares the resulting behavioral classification to human annotations. B_1,2_. We optimized our freezing classifier for analysis of rat and mouse freezing in videos obtained from other laboratories. C_1,2_. The Classifier Optimization Module iteratively sweeps through a range of thresholds for different metrics and reports F1 scores. The module first swept through a range of values for head/rear back velocity and change in head angle. D_1,2_. The highest F1 score (red box) was selected and then a subsequent sweep through two additional value ranges (for window width and count threshold from the smoothing algorithm) produced an equivalent or higher F1 score (red box). E. Once the user has identified candidate threshold values, the Validation Module reports recall, precision, F1, and specificity scores. F. The BehaviorDEPOT classifier performed robustly on videos of rats recorded in a different lab (Precision = 0.93; Recall = 0.88; F1=0.91; Specificity = 0.96). BehaviorDEPOT freezing classification was indistinguishable from highly trained human raters (N=4 videos, *P*=0.89, Mann-Whitney U). G. The BehaviorDEPOT classifier performed robustly on videos of mice recorded in a different lab (Precision = 0.98; Recall = 0.92; F1=0.95; Specificity = 0.95). BehaviorDEPOT freezing classification was indistinguishable from highly trained human raters (N=4 videos, *P*=0.49, Mann-Whitney U).

### The Validation Module

The Validation Module is designed to quickly assess a classifier’s predictive quality. To ensure that behavioral classifications are robust to different video recording conditions, the Validation Module compares classifier output to trained human raters across frames from multiple videos. The user will initially be prompted to indicate which classifier to evaluate and select a directory containing behavior videos and accompanying BehaviorDEPOT-analyzed tracking data. For each video, the module will categorize each frame as true positive, false positive, true negative, or false negative, using the human data as a reference. Precision, recall, specificity, and F1 score are then calculated and visualized for each video. These statistics are also reported for the total video set by concatenating all data and recalculating performance (Figure 7E). We used this data to validate the performance of the BehaviorDEPOT freezing classifier on the rat (Figure 7F) and mouse (Figure 7G) videos we acquired from external laboratories.

### Automated Quantification of Decision-Making Behaviors

BehaviorDEPOT can be flexibly adapted for behaviors beyond the realm of conditioned fear. Many labs study contingency-based, cost-benefit decision-making in mazes that involve choice points. For example, effort-based decision-making involves weighing whether it is worth exerting greater effort for a more valuable outcome. A common rodent assay for effort-based decision-making is the barrier T-maze task in which animals choose whether to climb over a barrier for a large reward (high effort, high value reward; HE/HVR) vs. taking an unimpeded path for a smaller reward (low effort, low value reward; LE/LVR)^30^. Well trained animals typically prefer HE/HVR choices, taking a direct route over the barrier with stereotyped trajectories. However, when reward or effort contingencies are adjusted animals demonstrate vicarious trial and error (VTE), thought to be a marker of deliberative (rather than habitual) decision-making. During VTE trials, animals may pause, look back-and-forth, or reverse an initial choice^31^. Several groups have identified neural signatures of VTE in hippocampus, striatum and prefrontal cortex suggesting that animals may be playing out potential choices (i.e. vicariously) based on an internal model of maze contingencies^31^. Simple, flexible, and automated analysis tools for detecting VTE and aligning this behavior with neural data would significantly enhance our understanding of deliberative decision-making.

We used BehaviorDEPOT to automatically detect VTE in a barrier T-maze task and to report the ultimate choice animals made during each effort-based decision-making trial. First, we trained a new DLC network to track mice as they navigated the T-maze. In one version of the task, mice decided whether to turn left to collect a small reward or to turn right and climb a wire mesh barrier for a large reward. We then changed the effort contingency by adding a second barrier to equalize effort across choices (Figure 8A,B). We used the BehaviorDEPOT Analysis Module to perform automated analysis of the task. We defined 8 regions of interest (Approach, Choice, Left Effort, Left Reward, Left Food Cup, Right Effort, Right Reward, and Right Food Cup). We then used the stored tracking data to automatically detect trials, which we defined as the first frame when the animal entered the approach zone until the first frame when the animal entered the reward zone, and to report the outcome of each trial (left choice or right choice) (Figure 8C,D).

**Figure 8.**
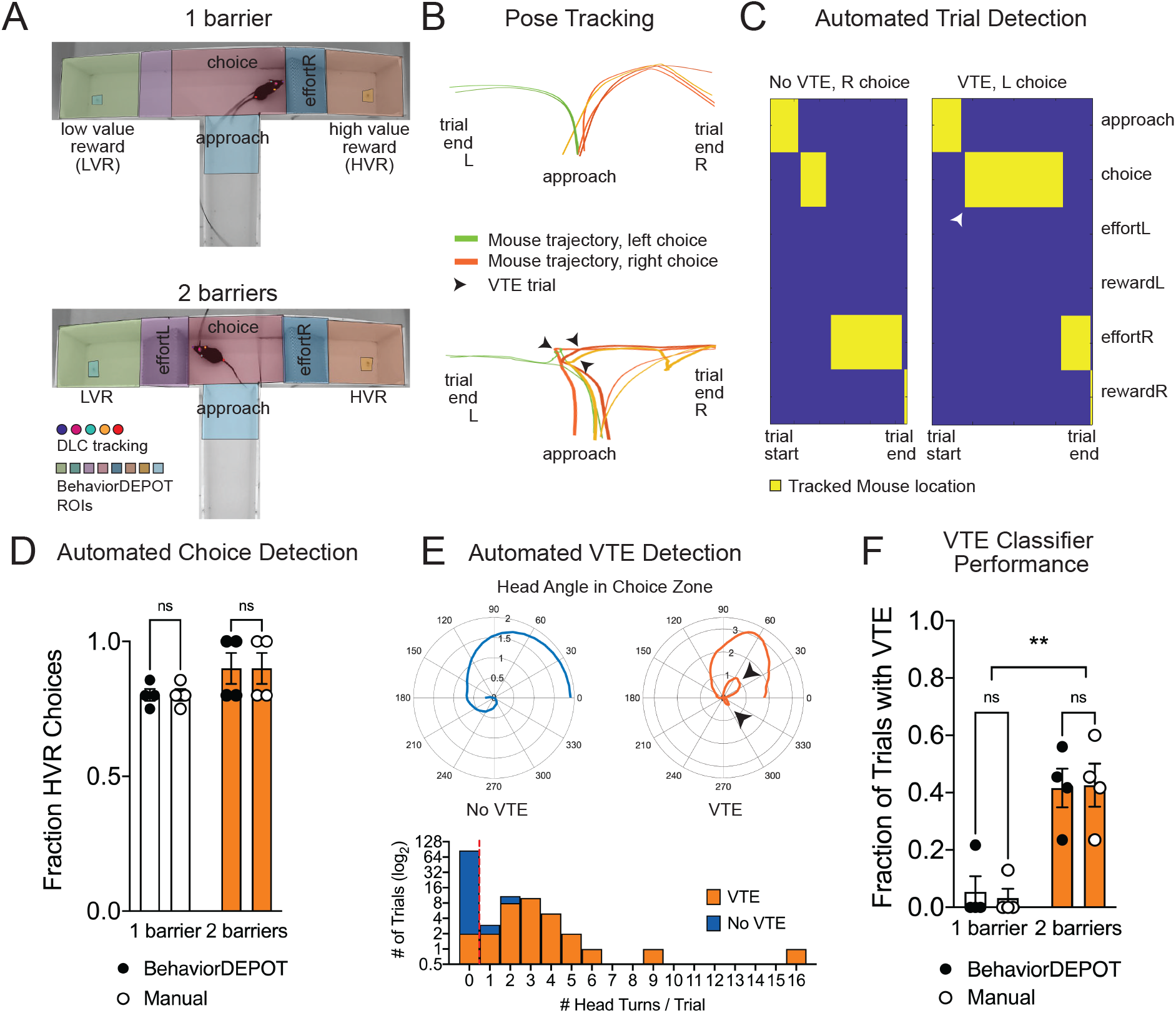
Use Case 4: Automated analysis of an effort-based decision-making T-maze task A. Screen shots showing DLC tracking in a 1-barrier (top) and 2-barrier (bottom) T-maze and ROIs used for analysis in BehaviorDEPOT. B. Sample mouse trajectories in a 1-barrier (left) and 2-barrier (right) T-maze. Lines represent individual trials for 1 mouse. Blue lines represent right choices, orange lines represent left choices, and thick lines indicate vicarious trial and error (VTE). C. Illustration of automated trial definitions. D. Automated choice detection using BehaviorDEPOT. BehaviorDEPOT indicated choice with 100% accuracy (*F*_*rater*_(1,6)=6.84, *P*>0.99, *F*_*Barriers*_(1,7)=4.02, *P*=0.09; *F*_*Subject*_(6,7)=0.42, *P*=0.84, 2-way ANOVA with Sidak post-hoc comparisons, 84 trials, N=4 mice). E. Top: Polar plots show representative head angle trajectories when the mouse was in the choice zone during a trial without VTE (left) and with VTE (right). Bottom: Histogram of head turns per trial for trials without VTE (blue) and with VTE (orange). Red dotted line indicates selected threshold. F. Fraction of trials with VTE during 1-barrier and 2-barrier sessions, comparing manual annotations to BehaviorDEPOT classification (*F*_*RaterxBarriers*_(1,6)=0.04, *P*=0.85, *F*_*rater*_(1,7)=0.03, *P*=0.85; *F*_*Barriers*_(1,6)=22.9, *P*=0.003, 2-way ANOVA with Sidak post-hoc comparisons, 102 trials, N=4 mice). Error bars represent S.E.M.

To develop a VTE classifier, we used the head angle data stored in the BehaviorDEPOT ‘Metrics’ data array. For each trial, we analyzed the head angles when the mouse was in the choice zone and used these values to determine the number of head turns per trial. Manually annotated VTE trials tended to have 1 or more head turns, while non-VTE trials tended to have 0 head turns, so we defined VTE trials as having 1 or more head turns in the choice zone (Figure 8E). We then used our BehaviorDEPOT VTE classifier to detect the fraction of trials with VTE in T-maze sessions with 1 vs. 2 barriers, finding a significant increase in the occurrence of VTE trials when a second barrier was added. Importantly, the BehaviorDEPOT performance was indistinguishable from a human rater (Figure 8E). These analyses highlight the utility of the robust repository of movement statistics that is automatically generated in BehaviorDEPOT. By calculating and storing information including velocity, angular velocity, head angle, and acceleration in a framewise manner, BehaviorDEPOT allows users to design automated analysis pipelines for a wide range of commonly studied cognitive tasks.

## Discussion

BehaviorDEPOT extends the utility of existing popular tools for markerless point tracking (e.g. DLC^3^ and LEAP^32^) to create an open source, flexible, reliable, and automated behavioral analysis pipeline. As optogenetics, miniscopes, and open-source technologies for extracellular recording become available to increasing numbers of labs, users with little coding experience need easy-to-use programs that can align detailed behavioral analyses with neurophysiological recordings and manipulations. BehaviorDEPOT offers a graphical interface-based pipeline for running experiments and classifying behaviors of interest with reference to spatial and/or temporal (e.g., tone times, optogenetic stimulation times) experimental considerations. Dedicated modules allow even inexperienced users to create, test, and optimize behavioral classifiers of their choosing.

We highlight the immediate utility of BehaviorDEPOT for analyzing behavior in widely used assays including contextual fear conditioning, active avoidance, and effort-based decision making. We report the first DLC-based freezing classifier with a user-friendly GUI and validate our freezing classifier in a variety of experimental designs with potential visual confounds: unoperated mice, mice and rats from our lab and others wearing an optogenetic patch cable, and mice wearing Miniscopes. In this way, our freezing classifier succeeds in scenarios in which commercially available alternatives often fail. To support miniscope-based analyses, we also report the development of the UCLA Minicam, which interfaces seamlessly with the BehaviorDEPOT to allow high quality tracking and automated behavior analysis in mice wearing UCLA Miniscopes. Using the classifier optimization module, we achieve robust freezing classification in videos of mice and rats recorded in external labs, highlighting the utility of BehaviorDEPOT for analyzing data that has already been collected. Finally, we leveraged the arrays of movement statistics that BehaviorDEPOT automatically generates to define trials in a T-maze task, report animal choice, and identify instances of VTE. These examples showcase BehaviorDEPOT as an easy-to-use, general utility software that is applicable to a wide range of behavioral neuroscience experiments.

BehaviorDEPOT fills an increasingly apparent gap in current resources. We integrate markerless point tracking with top-down recordings, enabling behavioral classification with two-dimensional spatial precision in an interface that requires no coding experience. Existing open-source software has succeeded in addressing other specific needs. For instance, ezTrack^9^ enables robust, Python-based freezing classification without the need of point tracking, but is limited to side-view videos, which constrains spatial analyses. MoSeq^6^ and DANNCE^8^ combine 3-dimensional video recordings with unsupervised machine learning to identify subsecond behavioral motifs, the complexity and coding-requirements of which may be beyond the scope of experimenters interested in more macro-level behaviors. Additional classifiers utilize multi-animal tracking for analyzing social behaviors^4,5^ and grooming behaviors^7,8^. With a well-validated freezing classifier and simple GUI that can run experiments and analyze, organize and align data, BehaviorDEPOT will bridge the gap between pose-tracking algorithms and neurophysiological recordings and manipulations for labs with any level of coding experience.

## Author Contributions

L.A.D., S.A.W., and C.J.G. conceived of experiments. C.J.G. and B.J. developed the freezing classifier. Z.Z. created MATLAB GUIs. C.J.G., B.J., Z.Z., and A.W. performed manual annotations of behavior. C.J.G. performed data acquisition and analysis of optogenetics experiments and Miniscope recordings. Z.Z. contributed to Miniscope data analysis. C.G. and D.A. designed and fabricated the UCLA MiniCAM and wrote accompanying software. M.J.S. and L.E.D. collected behavioral data from rats. C.J.G., Z.Z., B.J., S.A.W., and L.A.D. wrote the paper. All authors read and edited the paper.

## Acknowledgements

We thank A. Klein and N. Gogolla for contributing mouse behavioral videos for analysis with BehaviorDEPOT. We thank J. Reichl for assistance with DLC network training. This work was funded by NIH K01MH116264 (L.A.D.), Whitehall Foundation Research Grant (L.A.D), Klingenstein-Simons Foundation Grant (L.A.D.), NARSAD Young Investigator Award (L.A.D.), NIH T32MH073526 (B.J.), ARCS Pre-doctoral Fellowship (C.J.G.), NIMH K08 (S.A.W.).

## Methods

### Animals

Female and male C57B16/J mice (JAX Stock No. 000664) were group housed (2–5 per cage) and kept on a 12 hr light cycle. Following behavior conditioning, animals were individually housed until the memory retrieval sessions. All animal procedures followed animal care guidelines approved by the University of California, Los Angeles Chancellor’s Animal Research Committee.

### Contextual fear conditioning

Mice were handled for 5 days preceding the behavioral testing procedure. The conditioning chamber consisted of an 18cm x 18cm x 30cm cage with a grid floor wired to a scrambled shock generator (Lafayette Instruments) surrounded by a custom-built acoustic chamber. The chamber was scented with 50% Windex. Mice were placed in the chamber and then after a 2-minute baseline period, received 5 0.75mA footshocks spaced 1 minute apart. Mice were removed 1 minute after the last shock. Non-shocked control animals freely explored the conditioning chamber but never received any shocks. The following day, mice were returned to the conditioning chamber and a novel context (different metal floor, scented with 1% acetic acid), separated by a 1-hour interval. Context presentation order on day 2 was counterbalanced across mice.

### Platform-mediated avoidance

PMA used the fear conditioning chamber described above, except 25% of the floor was covered with a thin acrylic platform (3.5×4×0.5 inches). During training, mice were presented with 3 baseline 30s 4kHz tones (CS), followed by 9 presentations of the CS that co-terminated with a 2s footshock (0.13mA). The following day, mice were presented with 6 CS in the absence of shocks.

### Effort-based decision-making

For T-maze experiments, mice were food deprived to ~85% of their ad libitum initial weight (3-4 days) and habituated to handling and maze exploration. Reward pellets were chopped Reese’s peanut butter chips (~0.01g each). One arm of the maze was designated as high value reward (‘HVR’, 3 pellets), the other low value reward (‘LVR’, 1 pellet). Mice were trained to perform effort-reward decision-making via a sequential process. Once mice learned to choose the HVR arm (>80%), a 10 cm wire-mesh barrier was inserted to block that arm. Training was complete when mice achieved >70% success high effort/HVR choices on 2 consecutive days. Forced trials were used to encourage sampling of the barrier arm during training. Once mice achieved stable performance, a second barrier was inserted in the LVR arm to equalize effort between choices.

### Viruses

AAV1-syn-jGCaMP7f.WPRE (ItemID: 104488-AAV1) were purchased from Addgene and diluted to a working titer of 8.5×10^12^ GC/ml. and AAV1-CamKIIa-stGtACR2-FusionRed (ItemID: 105669-AAV1) were purchased from Addgene and diluted to a working titer of 9.5×10^11^ GC/ml.

### AAV injection with Optogenetic Cannula Implant

Adult wildtype C57/Bl6 mice were anesthetized with isoflurane and secured to a stereotaxic frame. Mice were placed on a heating blanket and artificial tears kept their eyes moist throughout the surgery. After exposing the skull, we drilled a burr hole above mPFC in both hemispheres (AP+1.8, ML+/−0.3 from bregma). A Hamilton syringe containing AAV-CaMKIIa-stGtACR2-WPRE was lowered into the burr hole and 400nL of AAV was pressure injected into each site (DV −2.25mm and −2.50mm from bregma) at 100nL/min using a microinjector (Kopf, 693A). The syringe was left in place for 10 minutes to ensure the AAV did not spill out of the target region. After injecting the AAV, chronic fiber-optic cannula (0.37NA, length = 2mm, diameter = 200um) were implanted bilaterally above the injection site and secured to the skull with Metabond (Parkell, S371, S396, S398). After recovery, animals were housed in a regular 12hr light/dark cycle with food and water ad libitum. Carprofen (5mg/kg) was administered both during surgery and for 2d after surgery together with amoxicillin (0.25 mg/mL) in the drinking water for 7d after surgery.

### Miniscope Surgery and Baseplating

For Miniscope recordings, all mice underwent two stereotaxic surgeries^21,33^. First, adult WT mice were anesthetized with isoflurane and secured to a stereotaxic frame. Mice were placed on a heating blanket and artificial tears kept their eyes moist throughout the surgery. After exposing the skull, a burr hole was drilled above PL in the left hemisphere (+1.85, −0.4, −2.3 mm from bregma). A Hamilton syringe containing AAV1-Syn-jGCaMP7f-WPRE was lowered into the burr hole and 400nL of AAV was pressure injected using a microinjector (Kopf, 693A). The syringe was left in place for 10 minutes to ensure the AAV did not spill out of the target region and then the skin was sutured. After recovery, animals were housed in a regular 12hr light/dark cycle with food and water ad libitum. Carprofen (5mg/kg) was administered both during surgery and for 2d after surgery together with amoxicillin (0.25 mg/mL) for 7d after surgery. Two weeks later, mice underwent a GRIN lens implantation surgery. After anesthetizing the animals with isoflurane (1–3%) and securing them to the stereotaxic frame, the cortical tissue above the targeted implant site was carefully aspirated using 27 gauge and 30-gauge blunt needles. Buffered ACSF was constantly applied throughout the aspiration to prevent tissue desiccation. The aspiration ceased after full termination of bleeding, at which point a GRIN lens (1mm diameter, 4mm length, Inscopix 1050-002176) was stereotaxically lowered to the targeted implant site (−2.0 mm dorsoventral from skull surface relative to bregma). Cyanoacrylate glue was used to affix the lens to the skull. Then, dental cement sealed and covered the exposed skull, and Kwik-Sil covered the exposed GRIN lens. Carprofen (5 mg/kg) and dexamethasone (0.2 mg/kg) were administered during surgery and for 7d after surgery together with amoxicillin (0.25 mg/mL) in the drinking water. 2 weeks after implantation, animals were anesthetized again with isoflurane (1–3%) and a Miniscope attached to an aluminum baseplate was placed on top of the GRIN lens. After searching the field of view for in-focus cells, the baseplate was cemented into place and the Miniscope was detached from the baseplate. A plastic cap was locked into the baseplate to protect the implant from debris.

### Optogenetics

Animals were habituated to the patch-cord for 3 days in advance of optogenetic stimulation. A patch-cord was connected to the fiber-optic cannula and animals were allowed to explore a clean cage for 5 minutes. On the testing day, optical stimulation through the fiber-optic connector was administered by delivering light through a patch-cord connected to a 473-nm laser (SLOC, BL473T8-100FC). Stimulation was delivered continuously with 2.5 mW power at the fiber tip.

### Miniscope Recordings

Mice were handled and habituated to the weight of the microscope for 4 days before behavioral acquisition. On the recording day, a V4 Miniscope was secured to the baseplate with a set screw and the mice were allowed to acclimate in their home cage for 5 minutes. Imaging through the Miniscope took place throughout the entire PMA training (~30 min) and retrieval session the following day. Behavior was simultaneously recorded with a UCLA MiniCAM.

### Behavior Video Recordings

Behavioral videos were acquired using one of the following 3 setups:

1. 50fps using a Chameleon3 3.2 megapixel monochrome USB camera fitted with a Sony 1/1.8 sensor (FLIR systems, CM3-U3-31S4M-CS) and a 1/1.8 lens with a 4.0-13mm variable focal length (Tamron, M118VM413IRCS). We recorded 8-bit videos with a 75% M-JPEG compression.
2. 30fps using a ELP 2.8-12mm Lens Varifocal Mini Box 1.3 megapixel USB Camera.
3. 50fps using a UCLA MiniCam 5 megapixel CMOS sensor (MT9P031I12STM-DP, ON Semiconductor)

### Histology

Mice were transcardially perfused with phosphate-buffered saline (PBS) followed by 4% paraformaldehyde (PFA) in PBS. Brains were dissected, post-fixed in 4% PFA for 12–24h and placed in 30% sucrose for 24–48 hours. They were then embedded in Optimum Cutting Temperature (OCT, Tissue Tek) and stored at −80°C until sectioning. 60um floating sections were collected into PBS. Sections were washed 3×10min in PBS and then blocked in 0.3% PBST containing 10% normal donkey serum (JacksonImmunoresearch, 17-000-121) for 2h. Sections were then stained with rabbit anti-RFP (Rockland 600-41-379 at 1:2000) in 0.3% PBST containing 3% donkey serum overnight at 4°C. The following day, sections were washed 3×5min in PBS and then stained with secondary antibody (JacksonImmunoresearch Cy3 donkey anti-rabbit IgG(H+L) 711-165-152, 1:1000) in 0.3% PBST containing 5% donkey serum for 2 hours at room temperature. Sections were then washed 5 min with PBS, 15 min with PBS+DAPI (Thermofisher Scientific, D1306, 1:4000), and then 5 min with PBS. Sections were mounted on glass slides using FluoroMount-G (ThermoFisher, 00-4958-02) and then imaged at 10x with a Leica slide scanning microscope (VT1200S).

### Manual annotation of behavior

Two-minute samples of each video recording were manually annotated by one-to-three highly trained individuals for freezing behavior. One-minute intervals were chosen from the beginning and end of the video recordings to capture diverse behaviors. In some cases, the entire video was annotated. Freezing was defined as the absence of movement except for respiration. Video frames (PointGrey: 36,000 frames; PointGrey+Opto: 36,000 frames; Webcam: 21,600 frames; MiniCAM: 46,069 frames; Rat videos: 40,320 frames; External mouse videos: 14,400 frames).

### Statistical analyses

Statistical analyses were performed in MATLAB or GraphPad Prism.

### Computer workstation specs

We trained networks in DLC and analyzed videos using two different custom-built workstations (Intel Core i9-9900K processor (8x 3.60GHz/16MB L3 Cache), 2×16GB DDR4-3000 RAM, NVIDIA GeForce RTX 2070 SUPER - 8GB GDDR6; AMD RYZEN 9 3950x processor (16×3.5GHz/64MB L3 Cache), 16GB DDR4 RAM, Gigabyte GeForce RTX 2060 SUPER 8GB WINDFORCE OC). BehaviorDEPOT can run on any personal computer and does not require a GPU.

### Installation of BehaviorDEPOT

Detailed instructions on BehaviorDEPOT installation can be found on GitHub: https://github.com/DeNardoLab/BehaviorDEPOT. Briefly, after installing a recent version of MATLAB (2018+), BehaviorDEPOT can be downloaded from Github and installed with a single click as either a MATLAB application or as a standalone exe file. Updates to the application will be added to the Github repository as they come available. We welcome feedback and bug reports on the BehaviorDEPOT Github page and encourage users to watch the page to be aware of any new releases.

### MiniCAM Instructions and Installation

Descriptions of fabrication and use of MinCAMs can be found on GitHub: https://github.com/Aharoni-Lab/MiniCAM.

---

**Supplemental Figure 1.**
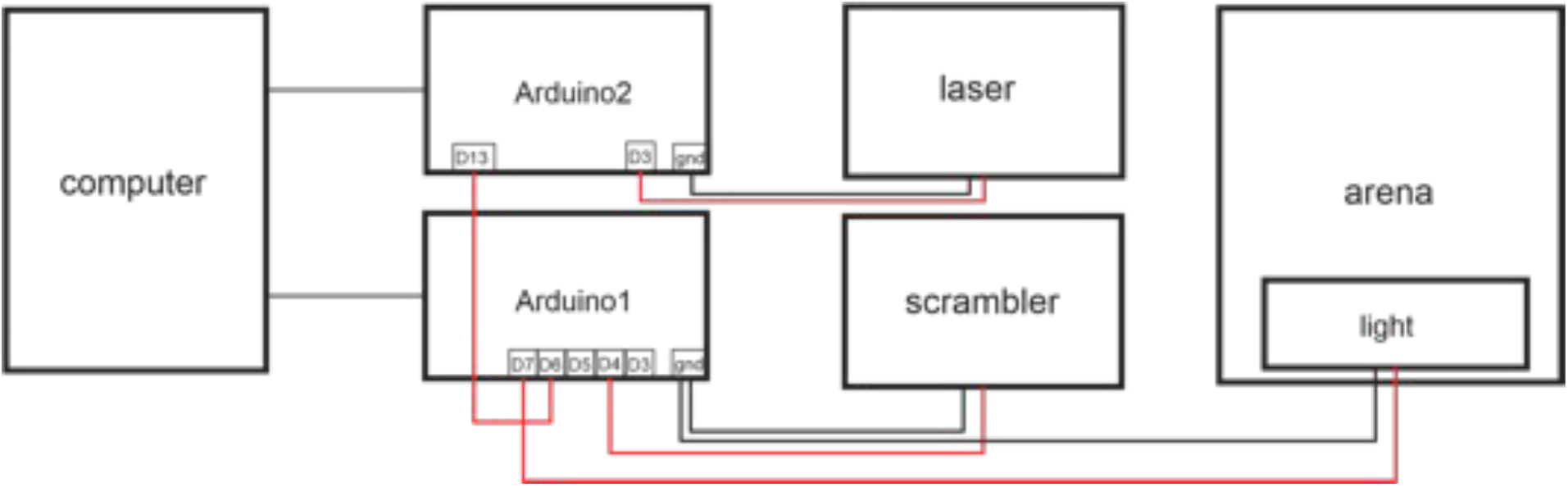
Example arrangement of Arduino interface between computer, fear conditioning and optogenetics hardware. The Experiment Module controls two Arduinos that control delivery of the scrambled shocker, and a light (for use as a conditioned cue), and the laser for optogenetics, respectively. MATLAB software triggers the conditioned tone.

**Supplemental Figure 2.**
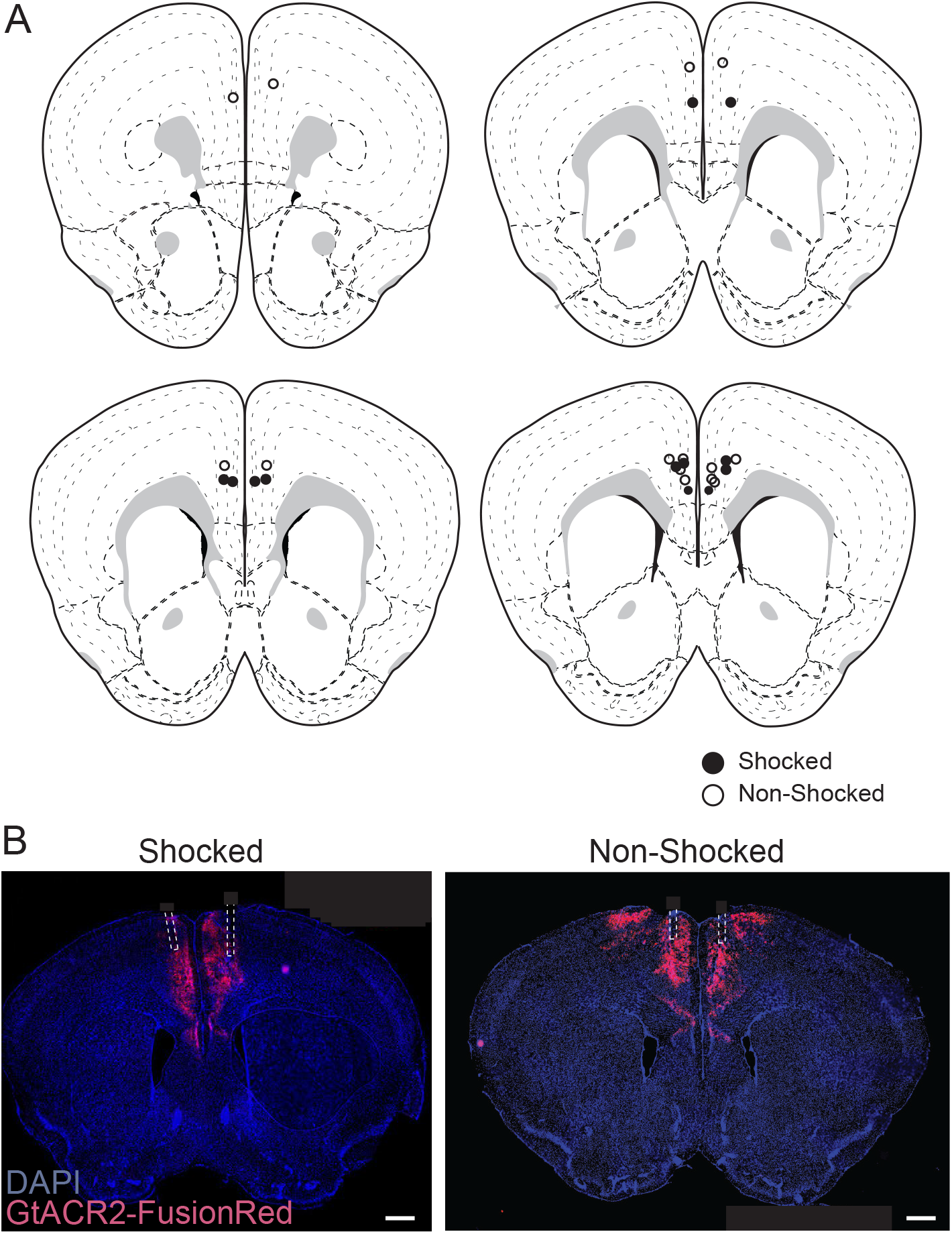
A. Optic fiber cannula placements for experiment described in Figure 3. B. stGtACR2 -FusionRed expression and bilateral fiber placement for representative shocked and non-shocked mice. Scale bar, 500um.

## References

1. Luo, L., Callaway, E. M. & Svoboda, K. Genetic Dissection of Neural Circuits: A Decade of Progress. Neuron (2018) doi:10.1016/j.neuron.2018.03.040.

2. Mathis, M. W. & Mathis, A. Deep learning tools for the measurement of animal behavior in neuroscience. Curr. Opin. Neurobiol. 60, 1–11 (2020).

3. Mathis, A. et al. DeepLabCut: markerless pose estimation of user-defined body parts with deep learning. Nat. Neurosci. 21, (2018).

4. Segalin, C. et al. The Mouse Action Recognition System (MARS): a software pipeline for automated analysis of social behaviors in mice. bioRxiv 2020.07.26.222299 (2020) doi:10.1101/2020.07.26.222299.

5. Nilsson, S. R. O. et al. Simple Behavioral Analysis (SimBA) – an open source toolkit for computer classification of complex social behaviors in experimental animals. bioRxiv 2020.04.19.049452 (2020) doi:10.1101/2020.04.19.049452.

6. Wiltschko, A. B. et al. Revealing the structure of pharmacobehavioral space through motion sequencing. Nat. Neurosci. 23, 1433–1443 (2020).

7. Hsu, A. I. & Yttri, E. A. B-SOiD: An Open Source Unsupervised Algorithm for Discovery of Spontaneous Behaviors. bioRxiv 770271 (2019) doi:10.1101/770271.

8. Dunn, T. W. et al. Geometric deep learning enables 3D kinematic profiling across species and environments. Nat. Methods 18, 564–573 (2021).

9. Pennington, Z. T. et al. ezTrack: An open-source video analysis pipeline for the investigation of animal behavior. Sci. Rep. 9, 1–11 (2019).

10. Hampel, F. R. The Influence Curve and its Role in Robust Estimation. J. Am. Stat. Assoc. 69, 383–393 (1974).

11. Cleveland, W. S. LOWESS: A Program for Smoothing Scatterplots by Robust Locally Weighted Regression. Am. Stat. 35, 54 (1981).

12. Corcoran, K. A. & Quirk, G. J. Activity in prelimbic cortex is necessary for the expression of learned, but not innate, fears. J. Neurosci. 27, 840–844 (2007).

13. Ledoux, J. E. Emotion circuits in the brain. Annu. Rev. Neurosci. 155–184 (2000).

14. Giustino, T. F. & Maren, S. The Role of the Medial Prefrontal Cortex in the Conditioning and Extinction of Fear. Front. Behav. Neurosci. 9, 1–20 (2015).

15. Sierra-Mercado, D., Padilla-Coreano, N. & Quirk, G. J. Dissociable Roles of Prelimbic and Infralimbic Cortices, Ventral Hippocampus, and Basolateral Amygdala in the Expression and Extinction of Conditioned Fear. Neuropsychopharmacology 36, 529–538 (2011).

16. Xu, W. & Südhof, T. C. A neural circuit for memory specificity and generalization. Science. 339, 1290–1295 (2013).

17. Pollack, G. A. et al. Cued fear memory generalization increases over time. Learn. Mem. 25, 298–308 (2018).

18. Frankland, P. W., Bontempi, B., Talkton, L. E., Kaczmarek, L. & Silva, A. J. The Involvement of the Anterior Cingulate Cortex in Remote Contextual Fear Memory. Science. 304, 881–883 (2004).

19. Mahn, M. et al. High-efficiency optogenetic silencing with soma-targeted anion-conducting channelrhodopsins. Nat. Commun. 9, 4125 (2018).

20. Ghosh, K. K. et al. Miniaturized integration of a fluorescence microscope. Nat. Methods 8, 871–878 (2011).

21. Cai, D. J. et al. A shared neural ensemble links distinct contextual memories encoded close in time. Nature (2016) doi:10.1038/nature17955.

22. Shuman, T. et al. Breakdown of spatial coding and interneuron synchronization in epileptic mice. Nat. Neurosci. 23, 229–238 (2020).

23. Dana, H. et al. High-performance calcium sensors for imaging activity in neuronal populations and microcompartments. Nat. Methods (2019) doi:10.1038/s41592-019-0435-6.

24. Barretto, R. P. J., Messerschmidt, B. & Schnitzer, M. J. In vivo fluorescence imaging with high-resolution microlenses. Nat. Methods 6, 511–512 (2009).

25. Jercog, D. et al. Dynamical prefrontal population coding during defensive behaviours. Nature(2021) doi:10.1038/s41586-021-03726-6.

26. Stout, J. J. & Griffin, A. L. Representations of On-Going Behavior and Future Actions During a Spatial Working Memory Task by a High Firing-Rate Population of Medial Prefrontal Cortex Neurons. Front. Behav. Neurosci. 14, 1–17 (2020).

27. Bravo-Rivera, C., Roman-Ortiz, C., Montesinos-Cartagena, M. & Quirk, G. J. Persistent active avoidance correlates with activity in prelimbic cortex and ventral striatum. Front. Behav. Neurosci. (2015) doi:10.3389/fnbeh.2015.00184.

28. Diehl, M. M. et al. Divergent projections of the prelimbic cortex bidirectionally regulate active avoidance. Elife 9, 1–13 (2020).

29. Lu, J. et al. MIN1PIPE: A Miniscope 1-Photon-Based Calcium Imaging Signal Extraction Pipeline. Cell Rep. (2018) doi:10.1016/j.celrep.2018.05.062.

30. Bailey, M. R., Simpson, E. H. & Balsam, P. D. Neural substrates underlying effort, time, and risk-based decision making in motivated behavior. Neurobiol. Learn. Mem. 133, 233–256 (2016).

31. Redish, A. D. Vicarious trial and error. Nat. Rev. Neurosci. 17, 147–159 (2016).

32. Pereira, T. D. et al. Fast animal pose estimation using deep neural networks. Nat. Methods 16, 117–125 (2019).

33. Shuman, T. et al. Breakdown of spatial coding and interneuron synchronization in epileptic mice. Nat. Neurosci. (2020) doi:10.1038/s41593-019-0559-0.

